# The interplay of structural and cellular biophysics controls clustering of multivalent molecules

**DOI:** 10.1101/373084

**Authors:** A. Chattaraj, M. Youngstrom, L. M. Loew

## Abstract

Dynamic molecular clusters are assembled through weak multivalent interactions and are platforms for cellular functions, especially receptor-mediated signaling. Clustering is also a prerequisite for liquid-liquid phase separation. But it is not well understood how molecular structure and cellular organization control clustering. Using coarse-grain kinetic Langevin dynamics, we performed computational experiments on a prototypical ternary system modeled after membrane-bound nephrin, the adaptor Nck1 and the actin nucleation promoting factor NWASP. Steady state cluster size distributions favored stoichiometries that optimized binding (stoichiometry matching), but still were quite broad. At high concentrations, the system can be driven beyond the saturation boundary such that cluster size is limited only by the number of available molecules. This behavior would be predictive of phase separation. Domains close to binding sites sterically inhibited clustering much less than terminal domains because the latter effectively restrict access to the cluster interior. Increased flexibility of interacting molecules diminished clustering by shielding binding sites within compact conformations. Membrane association of nephrin increased the cluster size distribution in a density-dependent manner. These properties provide insights into how molecular ensembles function to localize and amplify cell signaling.

## Introduction

Dynamic molecular clusters can form through weak binding interactions between multivalent molecules. They are highly plastic structures with a distribution of stoichiometries and sizes; they are becoming increasingly recognized as molecular platforms to drive key cellular functions, especially receptor-mediated signaling (2–12). For example, the epidermal growth factor receptor (EGFR) dimerizes and develops kinase activity when it binds its ligand, resulting in multiple phosphorylated sites on the cytoplasmic domains; these in turn interact with multiple SH2 domains on other multivalent scaffold or adaptor proteins, which then recruit additional binding partners (13). The result is the formation of multi-molecular dynamic ensembles with a distribution of stoichiometries and sizes, but with robust and specific cell signaling functions. We and others have used the term “ensemble” (4, 6, 7, 12, 14, 15) to specifically convey the notion of dynamic cluster composition and size. Formation of molecular ensembles is also a prerequisite for the phenomenon of liquid-liquid phase separation, which has become a major focus of cell biophysics research (16, 17).

The physical and chemical properties of molecular ensembles are not well understood, but it is clear that multivalency underlies their formation even when the individual binding affinities are weak. Biophysical models of these systems may serve to address the interplay of valency and geometry by systematically varying the cellular and molecular features underlying these properties. Stochastic reaction-diffusion modeling at the cellular scale cannot fully account for steric effects or molecular flexibility, because molecules are typically modeled as infinitesimal points in space (18–20). In principle, multimolecular/multistate interactions could be modeled with atomistic molecular dynamics simulations, but the large sizes of these systems and the need to simulate on the second timescale make such simulations computationally impractical.

In this work, we explore the interplay of multivalency and spatial effects with SpringSaLaD (21), a modeling and simulation software application developed in our lab that bridges the scales between molecular dynamics and cellular modeling. SpringSaLaD uses a Langevin dynamics formulation for linking spherical domains (or “sites”) with stiff springs to transmit the random diffusion-derived forces impinging on each site. It uses a unique exact formulation to relate bimolecular macroscopic on-rates to the microscopic reaction probabilities. Stochastic reaction-diffusion simulations are then run within a 3D rectangular spatial domain. We show how multivalency, membrane anchoring, steric interactions, molecular flexibility and the crowded cell environment all influence the size and distribution of molecular ensembles.

We do this by performing computational experiments on a prototypical multivalent system with a membrane anchored protein containing 3 phosphotyrosine (pTyr) sites, an adaptor protein containing one SH2 domain and 3 SH3 domains, and an effector protein consisting of 6 proline-rich motifs (PRMs) (22). This model is inspired by the nephrin - Nck1 - NWASP system, which we have previously studied with a non-spatial simulation algorithm based on Flory-Stockmayer theory (4). This system has also been extensively investigated experimentally by the Rosen lab (23, 24) and has been shown to exhibit liquid phase separation *in vitro.* Our model does not explicitly account for non-specific weak interactions between proteins such as electrostatics or the role of water in the liquid droplet phase. So we do not attempt to quantitatively model phase separation. However, we can show that at high concentrations cluster size is not self-limiting and grows as the number of available molecules increases, as would be expected for phase separation on a macroscopic scale.

## Methods

The SpringSaLaD simulation algorithm has been fully described (21), but will be briefly summarized here. Macromolecules are represented as a series of hard spherical sites that are linked by stiff springs. The motions of molecules are governed by a Langevin dynamics formulation that uses a diffusion coefficient assigned by the modeler to each site to calculate a distribution of random forces applied to each sphere. The forces are transmitted vectorially to neighboring sites in the molecule through the stiff spring linkers. Some of the spheres can represent binding sites, where the assigned on-rates with binding partners, together with their respective diffusion coefficients are used to calculate a reaction radius and reaction probability within each a time step. The shorter the time step the higher the accuracy of the simulations. A new bond is represented, simply, as a new stiff spring linking the binding sites. Inputs of off-rates are directly used to determine the probability of dissociation during a time step. In this study, we are primarily interested in characterizing the size and composition of clusters at steady state. We do this by initiating simulations with a random spatial distribution of elongated monomers and allowing them to diffuse and react stochastically until the average cluster size fluctuates around a stable size. We statistically analyze 50 stochastic trajectories for each condition. The 50 simulations are run in parallel on the CCAM High Performance Compute Cluster (https://health.uconn.edu/high-performance-computing/resources/); a typical run with 36 molecules for 500 ms requires 7 hours.

### Molecule construction

Our model has three molecular components – nephrin, Nck and NWASP. Each protein has multiple domains, which take part in biochemical interactions. While some of the specific domain structures are solved experimentally, the full protein structures are not yet available in the literature. So we have used secondary structure prediction homology modeling with web-server platforms like RaptorX (http://raptorx.uchicago.edu) (25) and Phyre2 (http://www.sbg.bio.ic.ac.uk/phyre2) (26) to generate approximate secondary structures from the amino acid (aa) sequences of interest. SpringSaLaD was then used to generate coarse grain models composed of multiple spheres linked with stiff spring linkers (Fig. 1). The relative sizes of the sites (Fig.S1) are determined by the predicted structures with the aid of a k-means clustering algorithm, *mol2sphere* (27).

**Figure 1:**
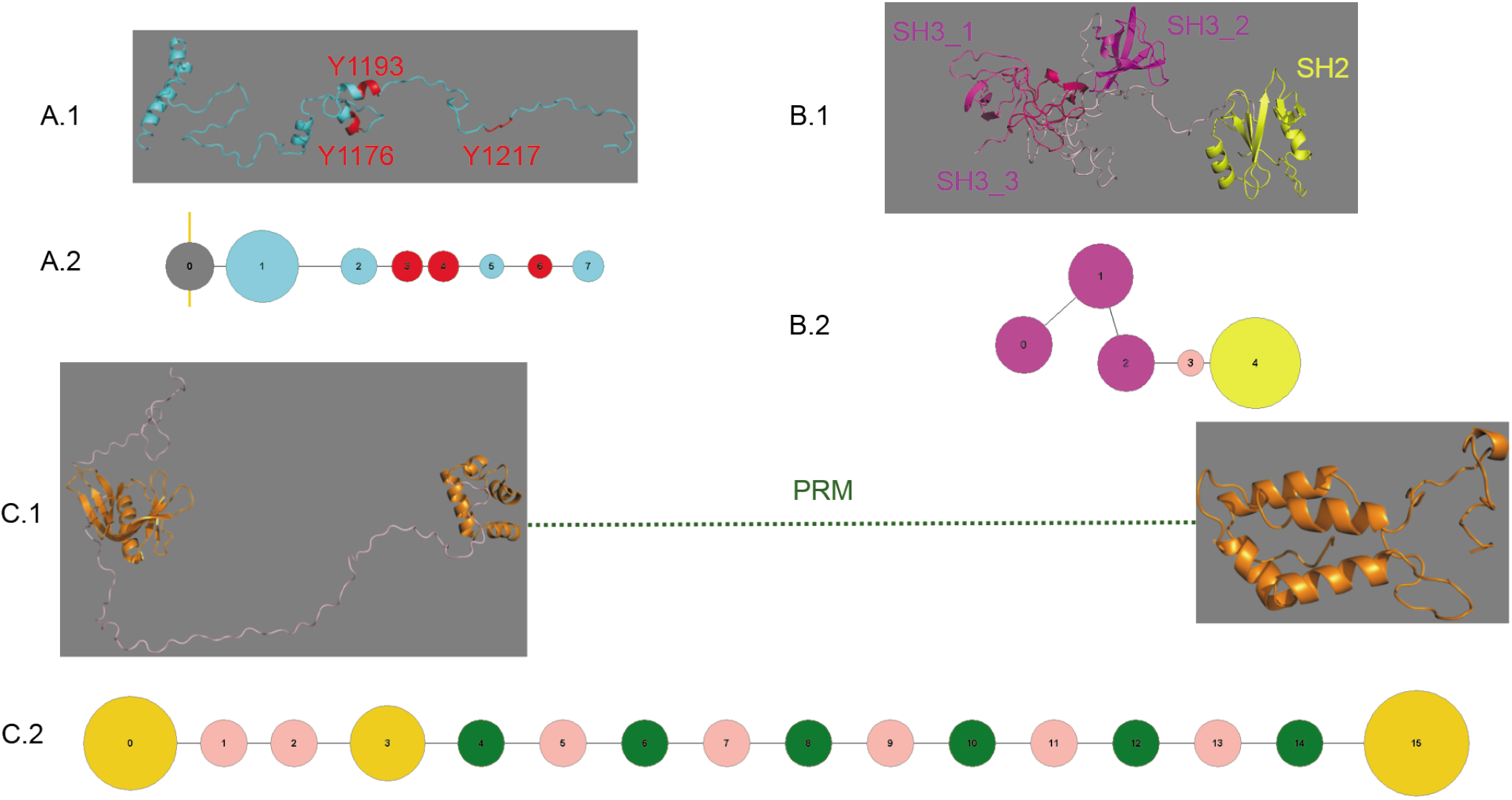
Construction of model components by coarse-graining the protein structures. (A) 165 aa long intracellular part of the transmembrane protein nephrin is modelled with 8 sites, three of them (red) represent pY-containing binding sites which bind to SH2 domains; this molecule is tethered with membrane (yellow linesurface) with the anchoring site (grey). (B) 376 aa long cytoplasmic protein Nck1 is modelled with three SH3 sites (magenta), one SH2 site (yellow) and one linker site (light pink).SH3_3 domain in the Nck1 secondary structure (B.1) is shown in reddish magenta for clarity. (C) 505 aa containing cytoplasmic protein NWASP has a lot of intrinsically disordered region (specially the proline rich motifs in 277-392) in its sequence. No secondary structure is predicted for the proline-rich sequence by most of the prediction software (shown in black dotted line). So this stretch is assumed to adopt polyproline-II (PP-II) type helical structure (average distance between two residues is 0.31 nm (1)) which is modelled with six binding sites (green) for SH3 domains and five structural sites (pink). The N-terminal (1–276) and C-terminal sequences (393505) are modelled with four and one structural sites respectively, according to the predicted secondary structures.

### Parameters

In SpringSaLaD, each site is assigned a diffusion coefficient (D). Because we are interested in the steady state cluster size distributions, as opposed to the kinetics of cluster formation, we could use somewhat smaller diffusion coefficients (Table 1) than realistic estimates; this permitted us to use longer time steps for our simulations, thus increasing computational throughput. Table 1 shows the D values assigned to the molecular sites. The high affinity binding of a pY site on nephrin to the SH2 domain on Nck is assigned a K_d_ of 1 μM (28); the low affinity binding between a Nck SH3 domain and a PRM on NWASP is assigned a Kd of 100 μM (29). Again, because the actual binding or dissociation rates do not affect the equilibrium cluster sizes, we chose values in Table 1 to optimize computational throughput. We checked some simulation results with both shorter timesteps and larger D values to assure that the steady state cluster characteristics were accurate when using our nominal values. All these input specifications can be found in the SpringSaLaD input files included in Supporting Information.

**TABLE 1:** PARAMETERS USED IN THE REFERENCE MODEL

|  |  |
| --- | --- |
| **[NEPHRIN]** | 9.06 $\mu\text{M}$ |
| **[NCK]** | 33.22 $\mu\text{M}$ |
| **[NWASP]** | 16.61 $\mu\text{M}$ |
| **KD** SH2 (NCK) – PY (NEPHRIN) | 1 $\mu\text{M}$ ($k_{\text{on}} = 100 \mu\text{M}^{-1} \text{s}^{-1}$; $k_{\text{off}} = 100 \text{s}^{-1}$) |
| **KD** SH3 (NCK) – PRM (NWASP) | 100 $\mu\text{M}$ ($k_{\text{on}} = 100 \mu\text{M}^{-1} \text{s}^{-1}$; $k_{\text{off}} = 10000 \text{s}^{-1}$) |
| **D** NEPHRIN_ANCHOR_SITE | 0.05 $\mu\text{m}^2/\text{s}$ |
| **D** NEPHRIN_CYTOSOLIC_SITES | 1 $\mu\text{m}^2/\text{s}$ |
| **D** NCK_SITES | 2 $\mu\text{m}^2/\text{s}$ |
| **D** NWASP_SITES | 2 $\mu\text{m}^2/\text{s}$ |

## Results

### Analysis of cluster size and composition

The baseline reference system (with which other configurations will be compared) consists of 36 molecules in a cubic domain 100 nm on each side as described in Figs. 1 and 2. The proportion of nephrin:Nck:NWASP was 6:20:10, respectively and chosen to approximate the optimal stoichiometry for binding site interactions; that is, in the reference system there are 18 pTyr sites interacting with 20 SH2 sites and 60 SH3 sites interacting with 60 PRM sites. Typically, we analyzed outputs from 50 trajectories. We can display the steady state outputs of these simulations as histograms, as illustrated in Fig. 3. In Fig. 3A, the abscissa corresponds to the average cluster size at steady state from a single simulation; this is calculated as the number of molecules (36 for the reference system) divided by the number of clusters. Thus, if a simulation produces 2 clusters each containing 18 molecules and another simulation produced 2 clusters containing 1 and 35 molecules, they would each yield the same average cluster size of 18. Since the number of clusters and the number of molecules are both small integers, only certain average cluster sizes are mathematically possible as can be seen from the sparse distribution in the histogram (Fig. 3A). To better examine how molecules are distributed over all possible cluster sizes, we can use an analysis displayed in Fig. 3B. Here, the abscissa is every possible cluster size from 1 to 36 molecules; the ordinate is the fraction of molecules populating these cluster sizes. For the scenario described above, a pair of 18mers would produce a single histogram entry at 18 on the abscissa with fraction of 1 on the ordinate; a configuration containing 1 monomer with 1 35mer would have an abscissa entry at 1 with a height of 0.028 and at 35 with a height of 0.972. For both of these histograms, we calculate weighted means, indicated by the dashed lines in Figs. 3A and B, that we term, respectively, Average Cluster Size (ACS) and Average Cluster Occupancy (ACO). For our 2 extreme scenarios, the ACS is 18 for both; the ACO for 2 18mers is 18 and for the combination of monomer and 35mer, ACO is 34.06. We can compute the ACS and ACO for each of the time points to generate averaged trajectories (Figs. 3C. and D.). The kinetics for both methods of assessing clustering are similar, reaching steady state by 100 ms. To produce the histograms in Figs. 3A and B, we use this 100 ms relaxation of the system as an interval over which to sample five time points (100 - 500 ms). As each time point corresponds to 50 runs, 250 independent data points are used to plot the steady state distributions in Fig.3A, 3B. Supplemental Figure S2 shows 5 representative individual trajectories over 500ms; the fluctuations within each of these show the system repeatedly accesses large clusters only to fall back to the smaller average steady state cluster sizes. To test whether there might be an additional slow component toward the steady state, we ran simulations out to 10 seconds (Supplementary Figure S3); there was no evidence of any longer term change.

**Figure 2:**
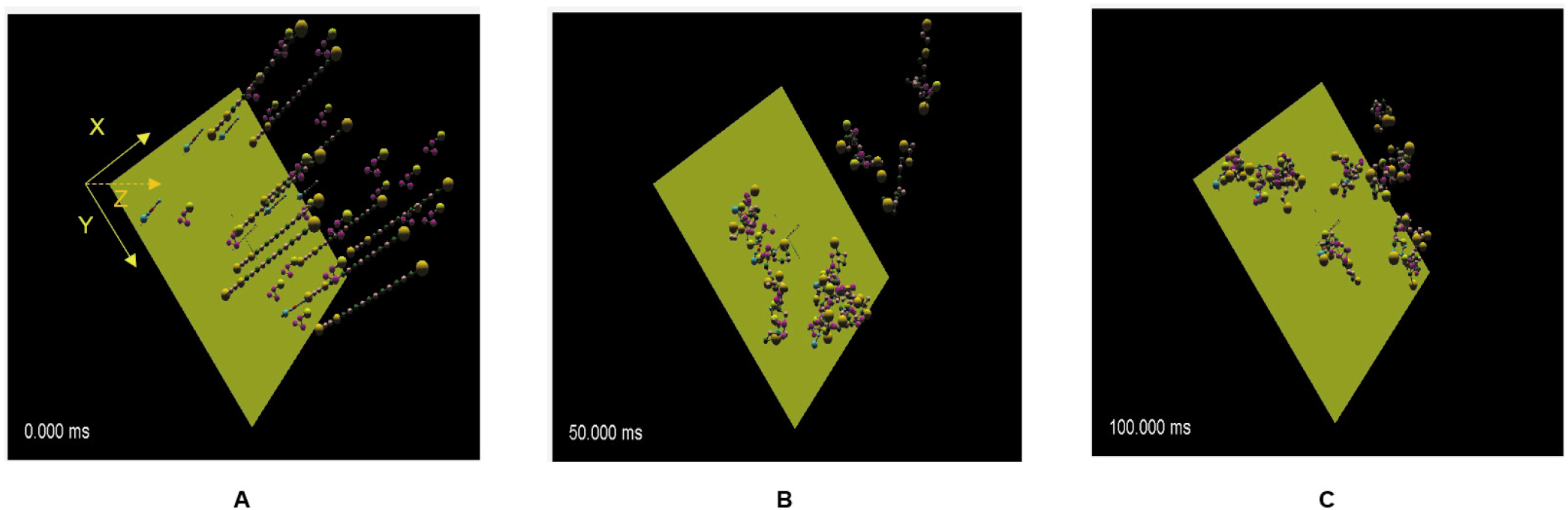
Clustering dynamics of the “reference” system. (A) The system contains a total of 36 molecules (6 nephrin, 20 Nck, 10 NWASP) in a cubic reaction volume (X = Y = Z = 100 nm) with 0 flux boundary conditions; XY surface represents the membrane where nephrin is anchored. At the beginning of simulation (t = 0), 36 molecules are distributed randomly (nephrin onto the membrane, Nck and NWASP in the cytosol) in the reaction volume. (B, C) Molecular clustering happens due to diffusion driven (multivalent) bimolecular interactions. At steady state (t ~ 100 ms), most clusters reside in the vicinity of membrane.

**Figure 3:**
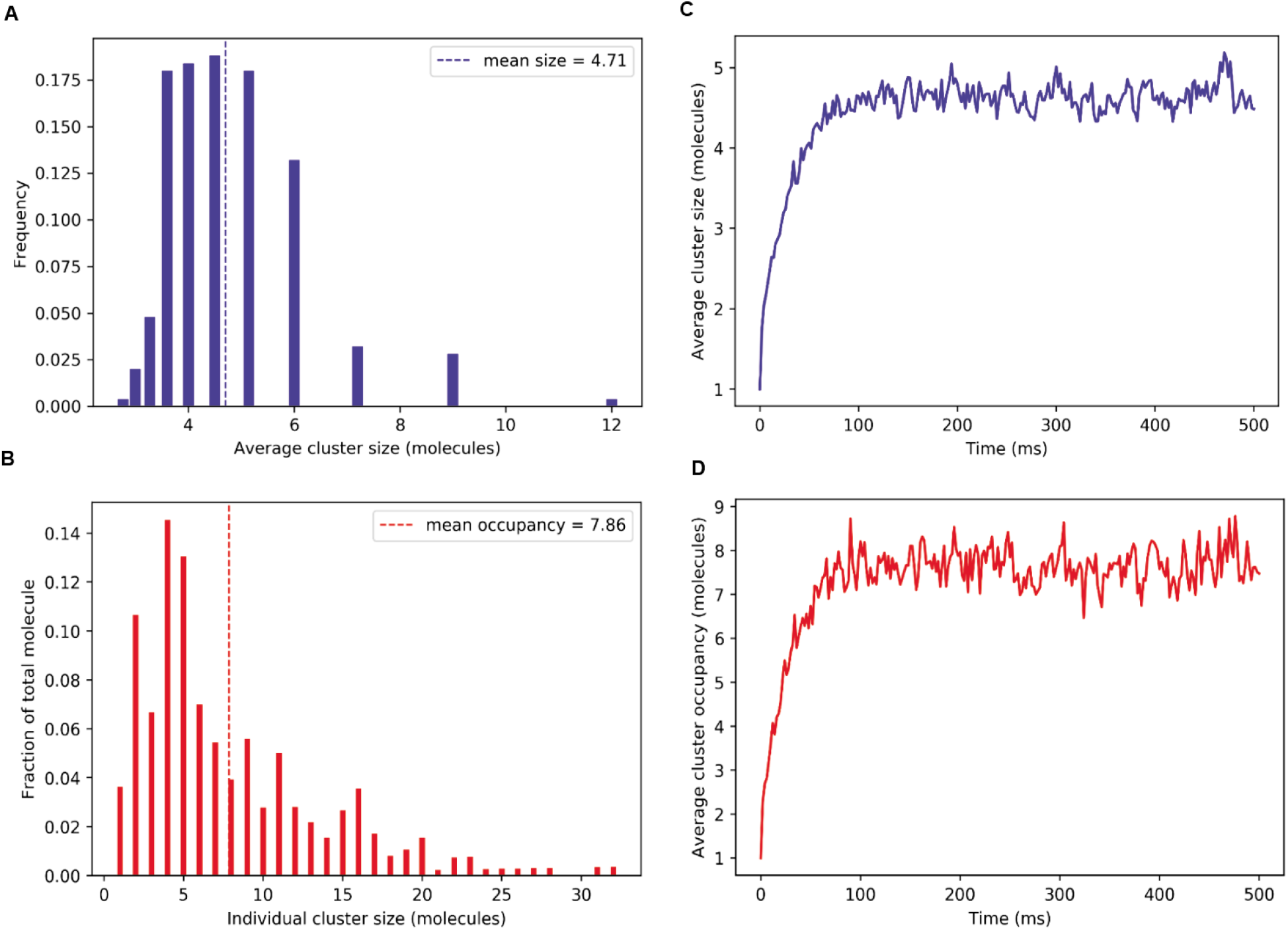
Quantification of molecular clustering. The steady state distributions (A, B) are sampled over five time points (100 - 500 ms) with the interval of 100 ms within all of the 500 trajectories. Thus, 250 independent samples are used to generate the histogram in A and B. Each time course (C, D) is averaged over 50 trajectories.

### Cluster size is insensitive to the number of available molecules until a concentration threshold is exceeded

A surprising property is the insensitivity of the ACS and ACO to the system’s size for this reference system. When we increase or decrease the number of molecules, keeping the molecular concentrations the same by adjusting the size of the domain, cluster size distributions and means turn out to be similar (Fig. 4). This indicates that the small number of available monomers does not limit the size of the clusters. Rather, this similarity in cluster sizes suggests that the system is sufficiently large to be approximated by equilibrium thermodynamics. In particular, there appears to be a balance of enthalpy and entropy that governs the size distributions of clusters and does not allow monomers to condense into a single large cluster. With a larger number of molecules, the average cluster size displays a much tighter distribution (Fig. 4A, bottom) because the higher number of possible average cluster sizes reduces the stochasticity of each average for individual steady state points. However, remarkably, there is very little effect on the shape of the distribution of fractional occupancy (Fig. 4B). This suggests that the shape of the distributions in Fig. 4B correctly reflects the tendency of the system to favor certain cluster sizes, as will be further discussed below. Overall, this analysis also indicates that our reference system with 36 molecules, is sufficiently large to provide a good representation of the steady state system behavior.

**Figure 4:**
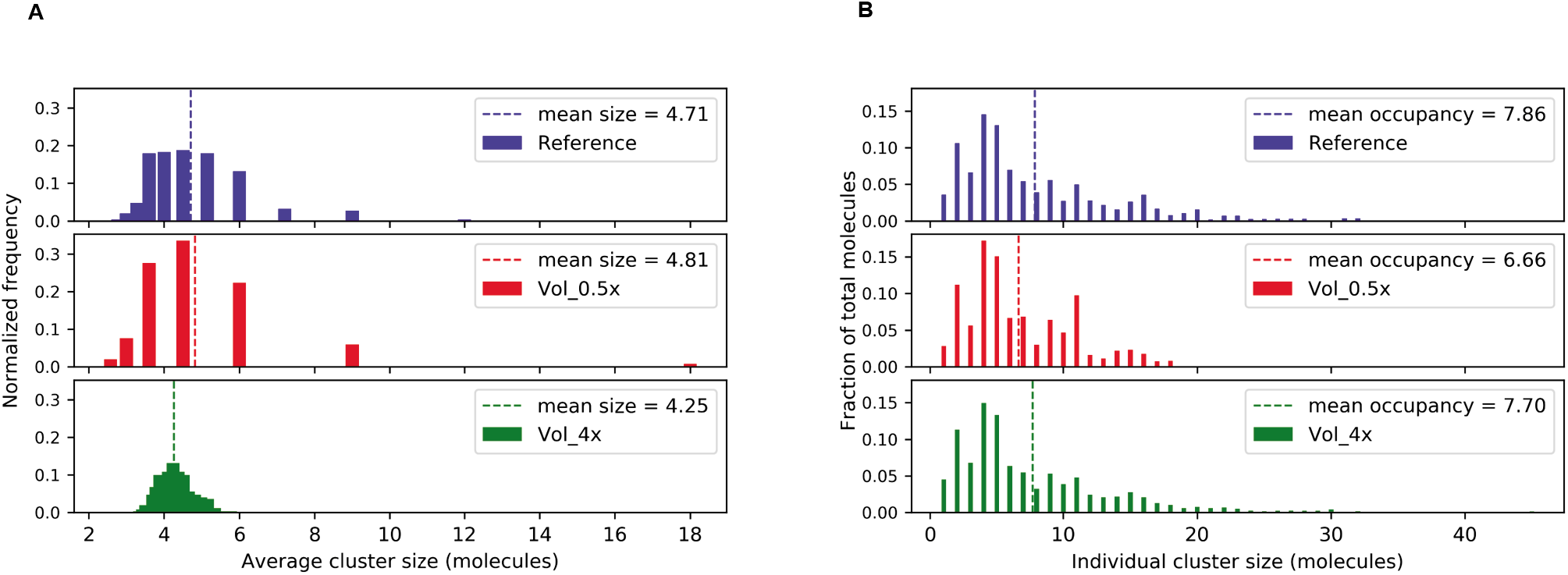
Clustering behavior is independent of system size for the reference system. (A, B) Steady state distribution of average and individual cluster sizes. The number of molecules are: 36 (blue), 18 (red) and 144 (green). The volume and membrane surface area is adjusted to maintain the same concentration and surface density in each simulation.

Increasing the affinity of binding (i.e. decreasing the Kds in Table 1), would be expected to produce larger clusters that may no longer be self-limiting as a function of system size. We tested this (Fig. 5) and found that the mean cluster size does increase compared to the reference system, as expected (compare Fig. 4), but the cluster size is still self-limiting. Interestingly, the histogram becomes quantized as strong intra-cluster interactions tend to saturate all the binding sites before clusters can grow. This behavior at high affinity, which is clearly a reflection of stoichiometry matching in these multivalent systems, has been previously compared to a “magic number” effect in physics (30), but is also akin to balancing a reaction equation in chemistry. For example, the prominent 11mer peak in the histograms in Fig. 5, corresponds to a nephrin_2_Nck_6_NWASP_3_ cluster - the smallest cluster in which each binding site is fully occupied. Even in the weaker binding system of Fig. 4B, the shape of the distribution of fractional occupancies remains remarkably the same as the system size increases. This reveals a tendency to maximize the binding site occupancy in individual molecules by favoring certain stoichiometries.

**Figure 5:**
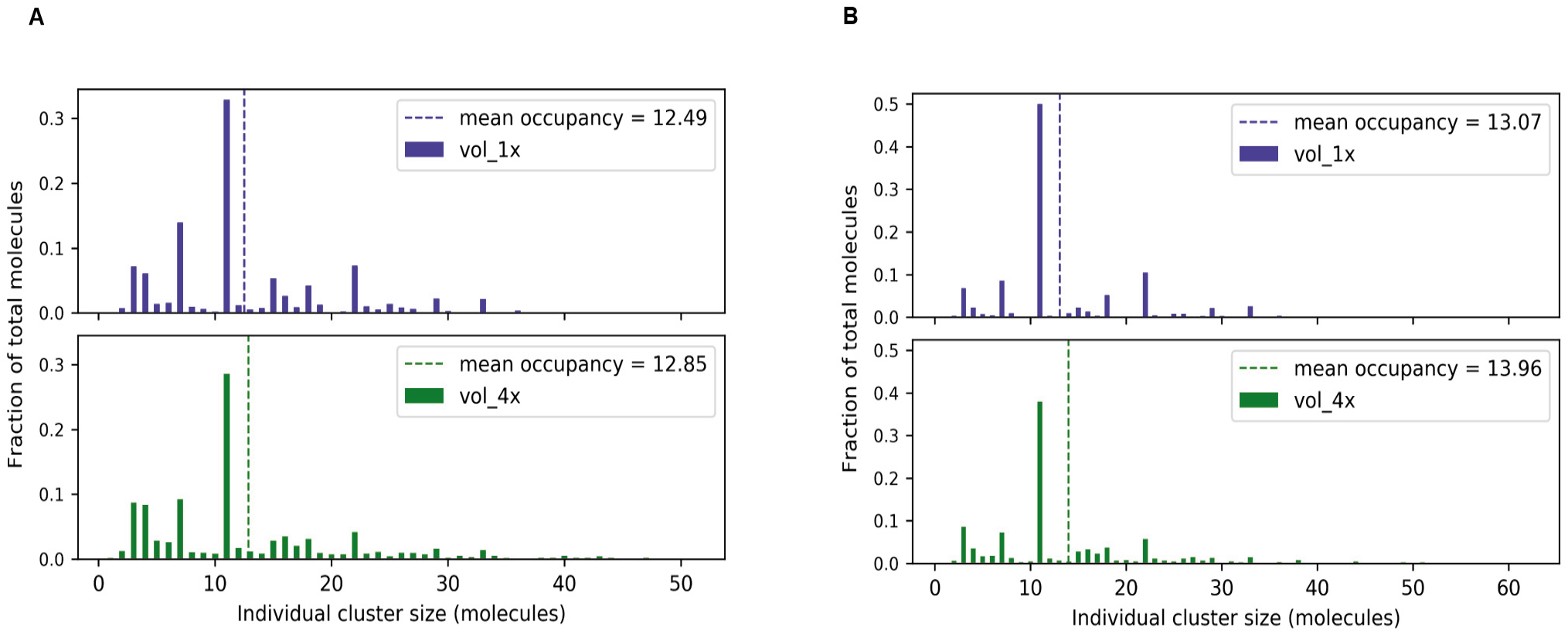
Higher affinity binding quantizes the cluster size distributions but the mean occupancy is still self-limiting. (A) Kds are reduced by a factor of 10 from those in Table 1; (B) Kds are reduced by a factor of 100 from those in Table 1. The upper row is for 36 molecules in a 100nm X 100 nm X100nm volume; the lower row is for 144 molecules in a 200nm X 200nm X 100nm volume.

We examined whether the cluster size was self-limiting at higher concentrations by raising the concentrations 2 fold and 4 fold (Fig. 6 and Fig. S4). For 2X concentrations, the system was still self-limiting; specifically, increasing the number of molecules from 72 (in a 100nm X 100nm X100nm volume) to 144 (in a 200nm X 100nm X100nm volume) produced the same distribution of cluster sizes with an average occupancy of ~15. However, for 4X concentration, the average occupancy continues to increase steadily up to the largest system size of 576 molecules (in a 200nm X 200nm X100nm volume) (Fig. 6, green plot). Thus, the 4X concentration has taken our system beyond the saturation boundary and cluster size distributions are no longer self-limiting. This system, which displays cluster distribution with a long tale and some clusters larger than 400 molecules (Fig. S4), would be predicted to display macro-aggregates corresponding to a phase separation.

**Figure 6:**
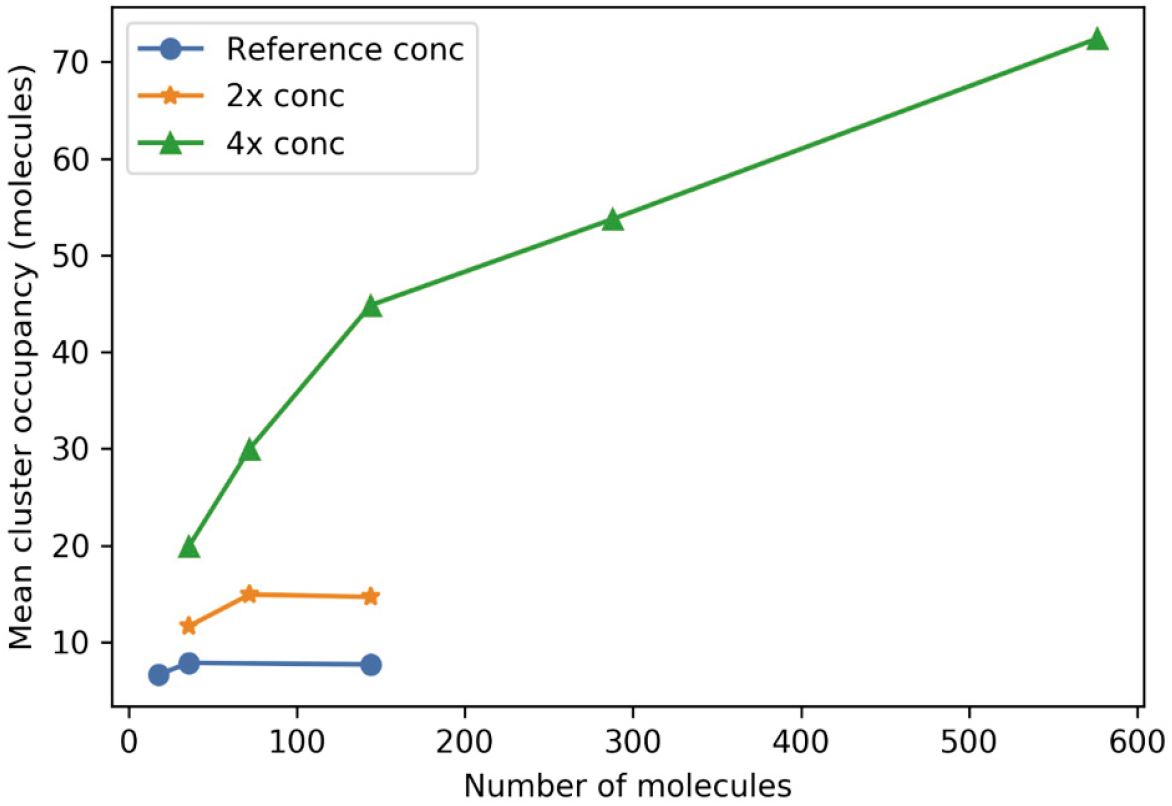
At high enough concentrations, the cluster size is not self-limiting. The concentrations are relative to the reference system (Table 1). Details of the dimensions of each simulation volume are given in Supporting Fig. S4, together with the full histogram corresponding to each of the points in this plot.

### Multivalency and optimal stoichiometry increases cluster size

Molecular clustering is a direct consequence of multivalent interactions between binding partners, but how this works in a specific system can be hard to predict. Here, we directly address this problem computationally. We manipulate the nephrin molecule to create three situations where the total number of binding sites are same, but the valency states are different (Fig.7A): 3 pY sites on each of 6 nephrin (3v, the “reference” system), 2 pY sites on each of 9 nephrin (2v), and 1 pY sites on each of 18 nephrin (1v). Nck and NWASP configurations are the same for all 3 scenarios. The histogram of fractional occupancy shifts to the left and, correspondingly, ACO decreases with reduced valence state (Fig. 7B). In considering this, it is important to appreciate that the strong interaction between the pY sites on nephrin and the single SH2 site on Nck assures that almost all the pY sites will be occupied with Nck in all 3 scenarios (as confirmed by the simulations results). Therefore, the effect of nephrin valency is actually to gather multiple Nck into a reduced local volume for interaction with the NWASP molecules. It should also be noted that this multivalency effect is over and above the effect of localizing the nephrin to the membrane, which actually becomes densely covered with nephrins for the 1v case, with 18 nephrins/10^4^nm^2^ (the effect of membrane localization is fully considered below).

**Figure 7:**
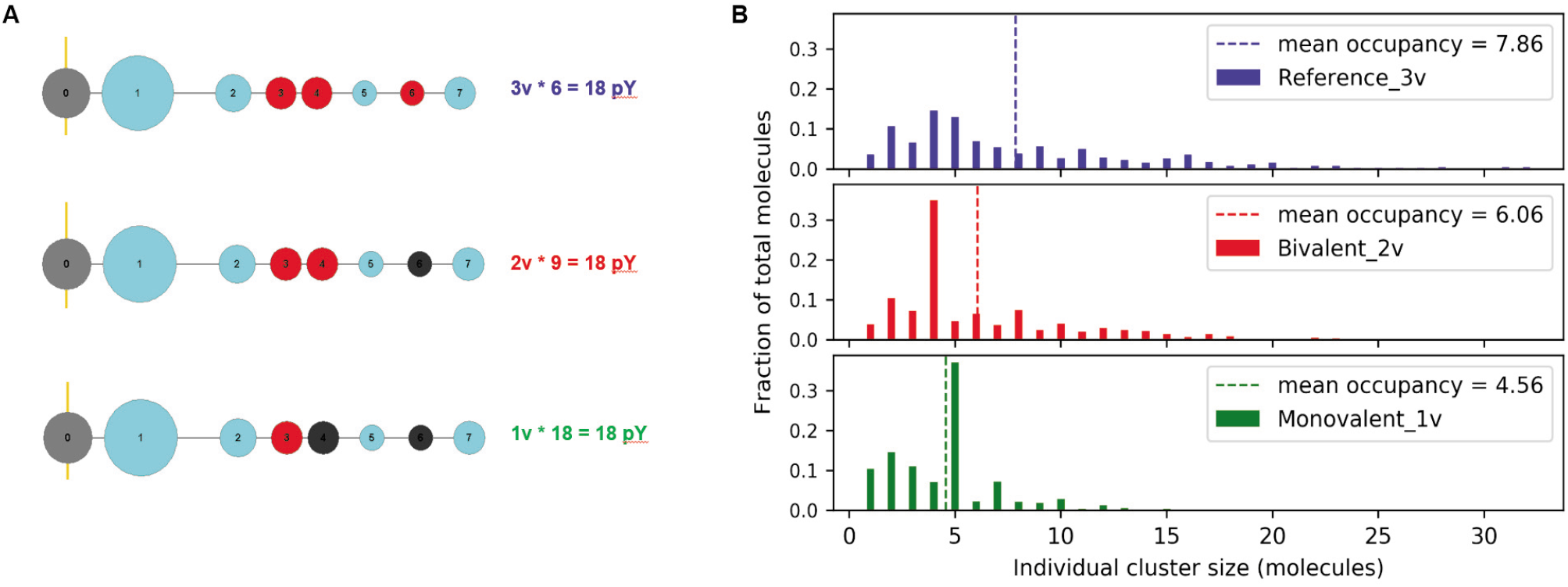
Multivalency increases cluster sizes. (A) Three Nephrin constructs with different valency; red sites represent pY (SH2 binding sites). The number of these tri, bi, and monovalent Nephrin molecules are adjusted (6, 9, 18 respectively) such that the total number of pY-sites remains same in all three cases. (B) Steady state distributions of individual cluster sizes (the dotted line represents the mean of the distribution).

Fig. 7B shows how the broad distribution of the reference trivalent nephrin system gives way to much sharper distributions when the nephrin valency is decreased. Strong preference for tetramers and pentamers are seen for the bivalent and monovalent, respectively, nephrin molecules. Again, optimal binding site stoichiometry underlies these preferences. The perfect binding site stoichiometry (nephrin: Nck: NWASP) for the bivalent system is 1:2:1 and for the monovalent nephrin, 2:2:1, accounting for the preference for tetramers and pentamers seen in Fig. 7B and Table S1. Overall, this analysis indicates that at a fixed affinity, increasing multivalency leads to larger clusters, but with broader size distributions.

As noted above, the count ratio of each molecule in the reference system, 6:20:10, was chosen to optimize the possible number of interactions between, respectively, nephrin, Nck and NWASP. We asked how clustering might be affected by altering the ratios of these molecules, keeping the total number of molecules at 36 (Table. S2). As expected, the reference system produces the largest clusters. Interestingly, however, a system where the ratio is, respectively, 10:20:6, is practically as good, despite the poor match between available binding site partners. What seems to be more important than the best match is the availability of a high level of Nck, which serves as an “adaptor” to link multivalent nephrin to multivalent NWASP.

### Interplay of steric effects, proximity effects and protein flexibility

Our model gives us a unique opportunity to probe the effect of molecular structural features on cluster formation. We first asked if the clustering propensity depends on the intramolecular distances between binding sites. We created two systems where the linker lengths within a molecule are elongated by 1 nm and 2 nm respectively compared with the reference system (Fig. 8A). As the distance between sites is increased, there is a dramatic shift in the distribution of molecules into larger clusters and a corresponding increase in mean cluster occupancy (Fig. 8B). The probable reason for this behavior is that with longer linkers, binding sites in the interior of a large cluster remain more accessible to additional binding partners; i.e. steric hindrance within the interior of a cluster is reduced with longer distances between binding sites.

**Figure 8:**
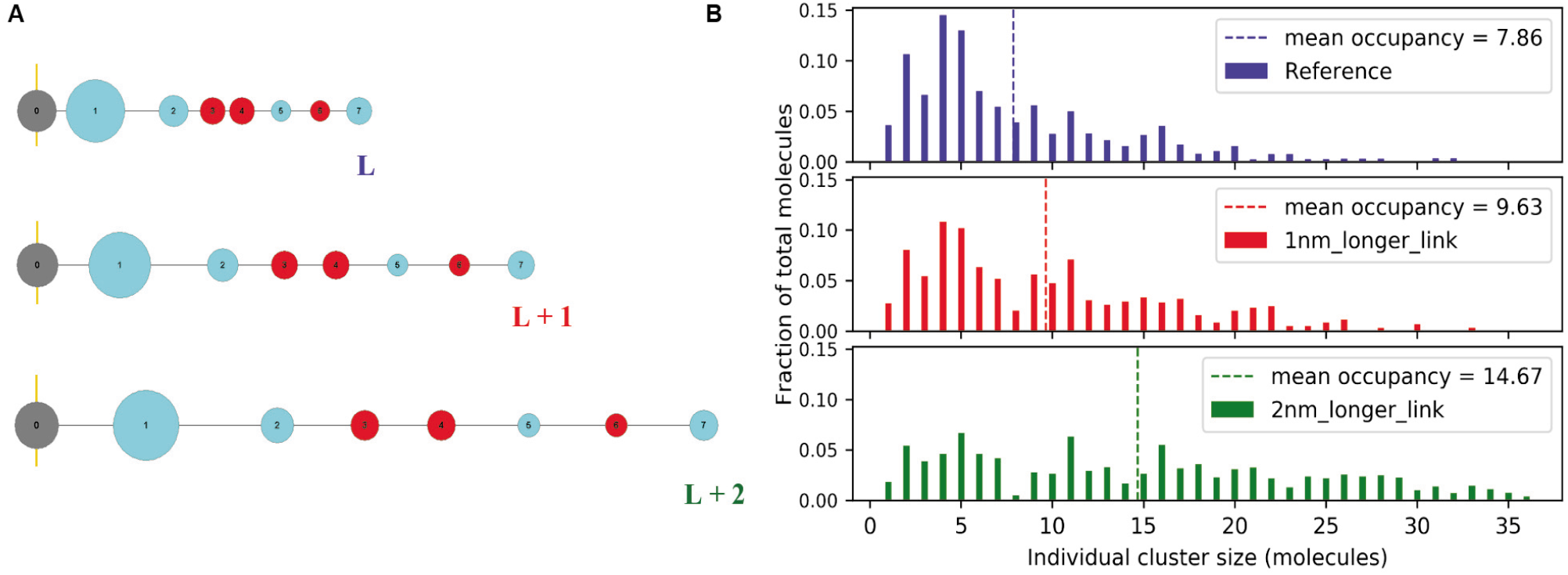
Effect of linker length. (A) The reference system compared to molecules where the linkers are increased by 1nm and 2nm. Only Nephrin is shown but the linker lengths were incremented in the same way in Nck and NWASp. (B) Steady state distributions of molecular occupancies (dotted line indicates the mean of the distributions).

While the analysis in Figure 8 is informative, in that it isolates the effect of distance between sites, the linkers in SpringSaLaD have 2 attributes that limit how they can represent the regions between binding sites: they are inflexible and they are sterically transparent. Therefore, to better represent the actual molecular structures, we included “structural” sites in all our molecules, as identified in Fig. 1. These do not have any binding attributes, but they do exclude volume and serve as pivot points for conformational flexibility. To isolate the steric effect, we create three NWASP constructs (Fig. 9) with similar flexibilities, but different steric effects by varying the sizes (Fig. S5) of the structural sites (nephrin and Nck retain their reference structures in all these simulations). Compared to the reference system (Fig. 9A), increasing the size of all structural sites (Fig. 9B) moderately decreases the clustering, while decreasing their sizes (Fig. 9C) moderately increases the cluster sizes. As would be expected, these changes reveal the influence of steric hindrance for binding interactions. Interestingly, removal of the peripheral structural sites from the reference NWASP molecule (Fig. 9D) dramatically increases the tendency to form larger clusters. This large effect of the peripheral binding sites can be attributed to their exclusion from the cross-linked interior of larger clusters (Fig. S6). This exclusion has the synergistic effects of unfavorably lowering the entropy of larger clusters and also serving as a steric barrier for the binding of monomers to available free binding sites in the interior. The 3 proteins that inspire this work all have intrinsically disordered region (IDR) in their sequences; the proline rich sequence in NWASP (31), the linker region between SH2 domain and first SH3 domain in Nck (24), and almost all the intracellular portions of the nephrin sequence (32). As a consequence of the flexibility of IDRs, the intramolecular distance between sites that flank them are more likely to vary in the course of interaction. To test the effect of flexibility, we created the NWASP molecules shown in Fig. 10 and simulated their clustering behavior with maintaining the reference structures of nephrin and Nck (Fig. 10A is the reference structure of NWASP). Interestingly, we see that the less flexible structure of Figs. 10B have the tendency to form very large clusters; structures with similar flexibilities but differently sized pivot sites (Fig. 10A vs. Fig. 10C) yield similar clustering behavior. This flexibility dependence could be explained by considering entropic effects. Since average cluster size is an equilibrium property, the loss of entropy would prevent the system from forming larger clusters, as discussed above. But the loss of entropy is not just due to the formation of multimers from monomers, but also because of the loss of conformational freedom resulting from the cross-linked multivalent binding. The more flexible the monomers, the greater the entropic loss upon cluster formation. Therefore, the steady state cluster size would go up with the less flexible initial structures, consistent with the trend shown in Fig. 10. Another way of explaining this would be the tendency of the less flexible molecules to populate relatively more extended conformations, where their binding sites remain more highly accessible. Turning to Fig. 10D, we see that decreasing both steric hindrance and flexibility produces the most dramatic enhancement of clustering, over and above the effects of each alone (compare Figs. 9D and 10B). Although we did not increase the system size (as in Fig. 6) to test if the cluster size distribution in Fig. 10D is self-limiting, we expect it is not and may be above the saturation boundary.

**Figure 9:**
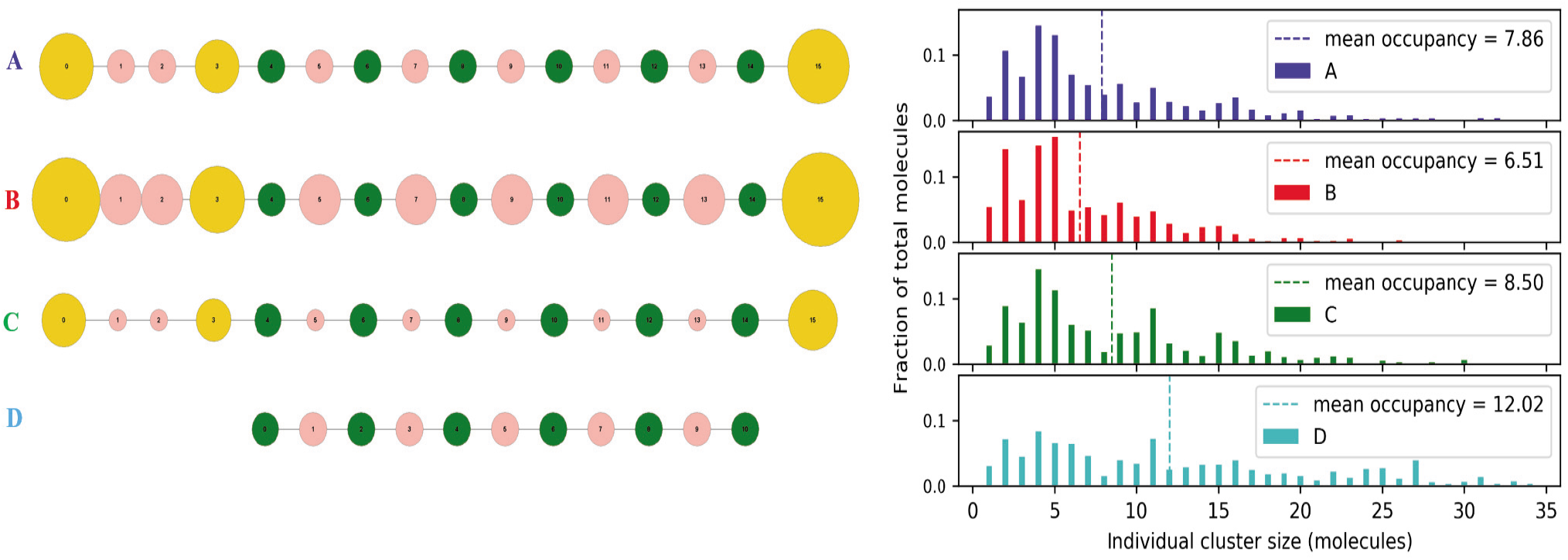
Non-binding structural sites in NWASP exert steric effects that reduce cluster size. Sizes of NWASP structural sites are varied, keeping the sizes of PRM binding sites (green) unchanged. (A) Reference; (B) Larger structural sites; (C) Smaller structural sites; (D) NWASP without the peripheral structural sites. Corresponding steady state distributions of individual cluster sizes (dotted line indicates mean of the distribution) are shown on the right. Only NWASP is varied in these computational experiments; nephrin and Nck retain their reference structures.

**Figure 10:**
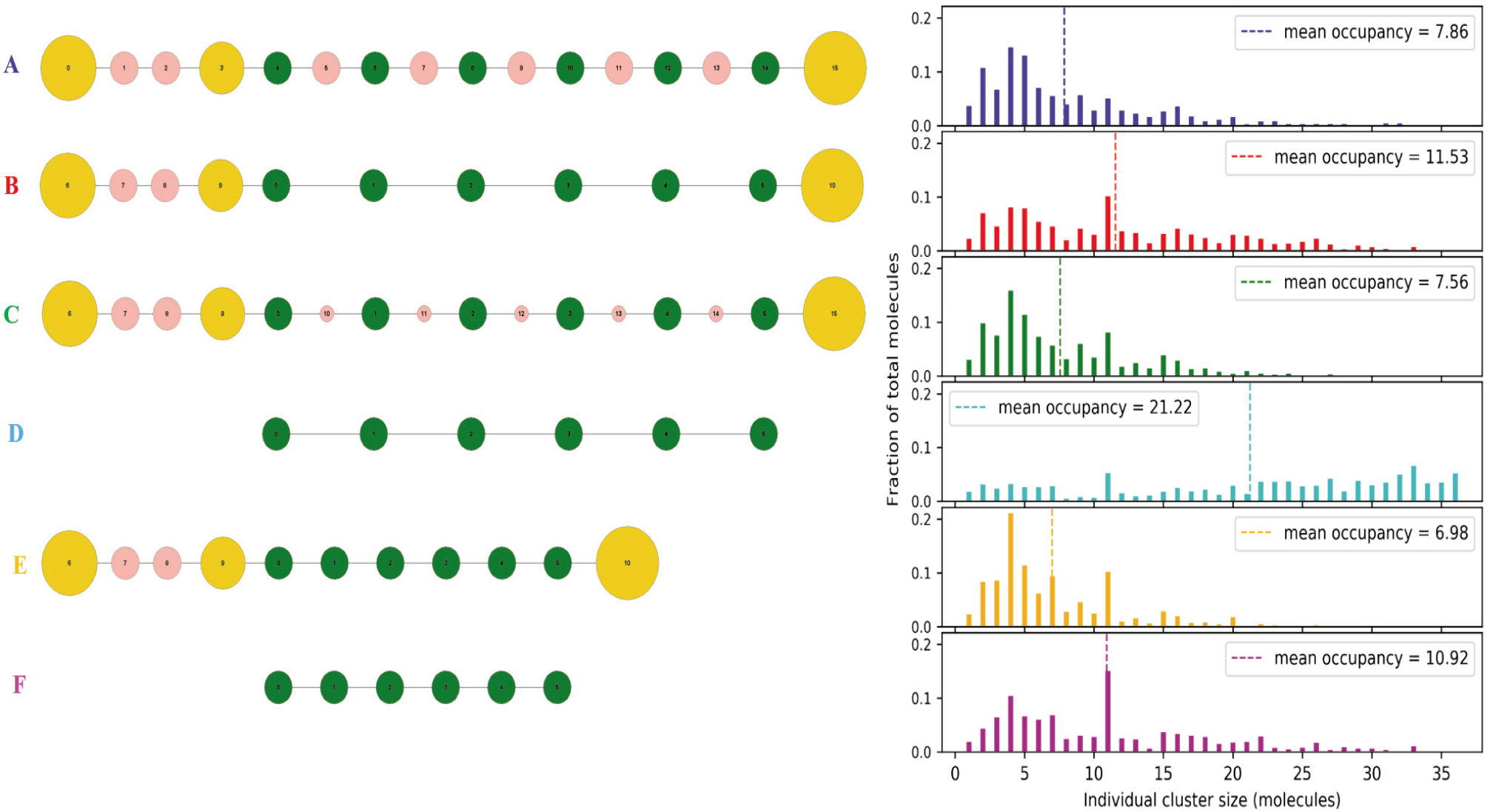
Flexibility within the binding domains and spacing between PRM sites control cluster sizes. (A) Reference configuration, (B) pivot sites (pink) flanking the PRM sites (green) are removed; (C) Similar to reference system, except the sizes of pivot sites in between PRM are smaller (diameter = 1 nm); (D) Bare PRM sites; (E) Similar to B except the linker lengths between PRM sites are shortened to 4 nm; (F) Bare PRM sites with 4 nm linker lengths. Nephrin and Nck configurations are kept same in all these cases. At the right are the corresponding steady state distributions of individual cluster sizes (dotted line indicates the mean of the distribution).

Figs. 10E and 10F point to the importance of spacing between binding sites, which may also be contributing to the trends in Figs. 10A-D. The PRM sites bind to SH3 sites in NcK. The latter are approximately 4nm apart (Fig. S1). In the NWASP structures of Figs. 10E and 10F, the distances between the PRM was reduced to 4nm compared to 7nm in the similar structure of Figs. 10B and 10D, repsectively. Reducing the distances between the PRMs to the distances between their SH3 binding partners promotes multiple bonds between pairs of molecules in a ladder-like configuration. This limits the ability to form larger clusters by promoting the stoichiometry matching between arrays of complementary binding sites. Indeed, as discussed in connection with Fig. 5, the 11mer, which has the minimum number of molecules needed to occupy all binding sites, is prominent in the histograms for Figs. 10E and F. The flexibility of the PRMs around the pivot sites in Figs. 10A and C may also allow for this laddering effect to contribute to the smaller sizes of their clusters, in addition to the entropy contributions discussed above.

### Membrane localization promotes multivalent clustering

Signaling cluster formation takes place in the spatial region near the plasma membrane; so membrane associated proteins are likely to play a key role in this process. In principle, membrane anchoring can promote clustering by producing by confining binding sites to a region of approximately one molecular length from the membrane; this effectively increases the local concentration compared to sites that are free within the bulk volume. On the other hand, the membrane might serve as a steric barrier preventing the binding partners to approach the membrane-tethered molecular binding sites. To understand the interplay of these 2 opposing effects, we created three nephrin constructs where length of the anchor linker is increased gradually (keeping the cytosolic Nck and NWASP as in the reference system) to reduce the potential steric effect from the membrane; we also include simulations where nephrin is detached from the membrane and free to diffuse around the cytosol, which should demonstrate the role of spatial confinement or effective higher local concentration around the membrane (Fig. 11). There is a small increase in fractional occupancy as a function of anchor length, indicating the steric contribution from the membrane is minor. Removing the membrane anchor results in a significant reduction of mean cluster occupancy, indicating that membrane confinement can potentiate clustering.

**Figure 11:**
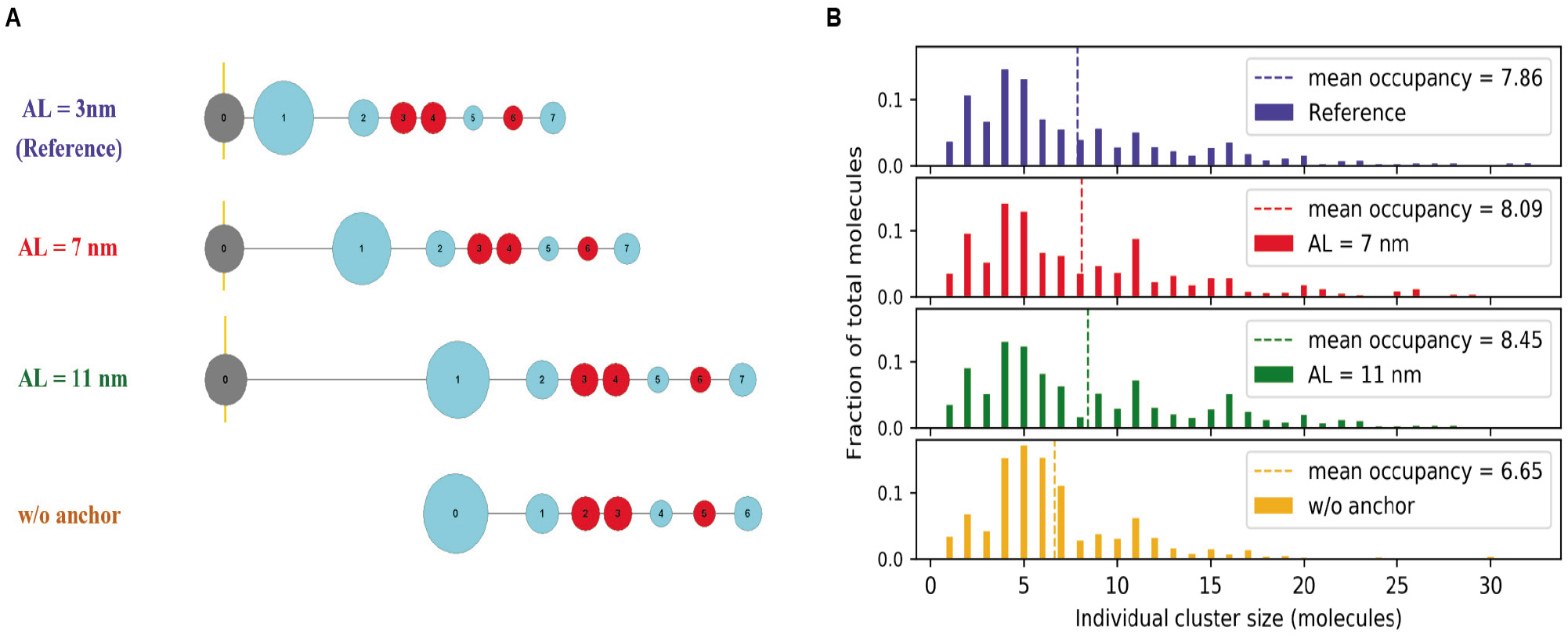
Membrane anchoring of nephrin promotes clustering. (A) Nephrin constructs with varying anchor length, AL (which is the linker that connects first structural site with the anchor site) and without anchor. (B) Steady state distributions of individual cluster sizes (dotted line indicates the mean of the distribution).

To further probe the importance of spatial confinement at the membrane, we explore the effect of density of nephrin molecules at the membrane surface. In our reference configuration, we have a total of 36 molecules (6 nephrin, 20 Nck, 10 NWASP) in a cubic (X = Y = Z = 100 nm) reaction volume with nephrin anchored on the XY plane. We can change the membrane surface density of nephrin, while maintaining constant volume and monomer concentrations, by altering the X, Y, Z dimensions (Fig. 12A, 12B, and 12C). When we increase the membrane density of nephrin in this way, we see a signifcant increase in average cluster size (Fig. 10D, E). Interestingly, the kinetics for approach to steady state is much slower for the higher density systems. As a control to assure that these effect are not associated with the aspect ratio, we performed simulations with the cubic and rectangular boxes in Figures 12 A and C, but without anchoring nephrin to the membrane (Fig. S7); without membrane anchoring, there is no difference in cluster size or cluster occupancy. Thus, the local confinement afforded by membrane anchoring significantly increases the propensity for clustering while decreasing the rate of cluster growth.

**Figure 12:**
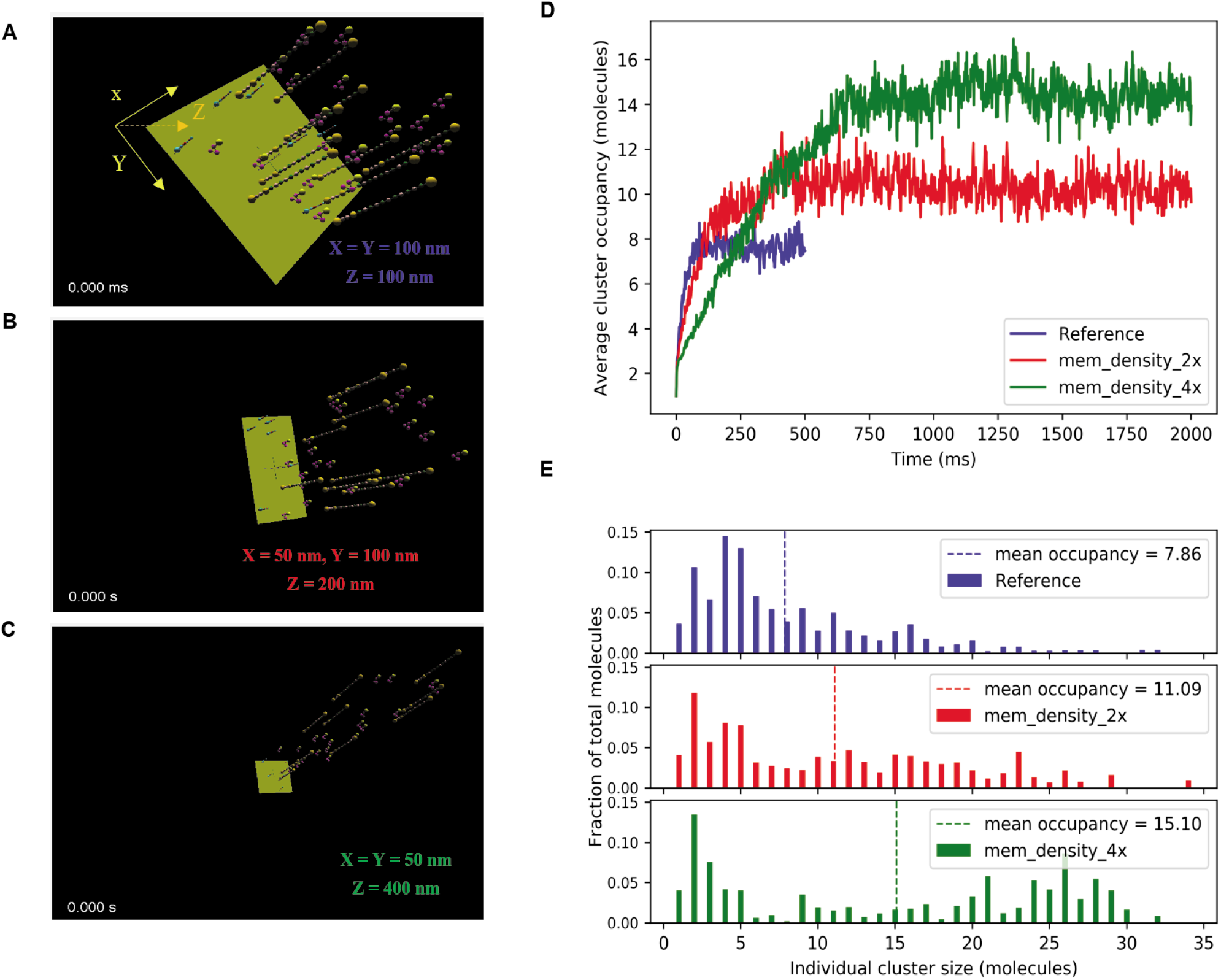
Increased nephrin membrane density promotes clustering. (A) Reference configuration in a cubic reaction volume (X = Y = Z = 100 nm); nephrin density = d (= 0.06 molecules.μm-2). (B) X = 50 nm, Y = 100 nm, Z = 200 nm; nephrin density = 2*d. (C) X = Y = 50 nm, Z = 400 nm; nephrin density = 4*d. (D) Dynamics of average molecular occupancy; note that times to reach steady states are different. (E) Steady state distributions of molecular occupancies. For higher nephrin density cases (red and green), two steady state time points (100 realizations) are sampled, unlike the reference system (250 realizations). Total number of molecules are the same in all simulations.

### Crowded cellular environment enhances clustering

Cells contain a wide range of biomolecules with varying shapes and sizes; these molecules occupy physical volume and behave as obstacles to other diffusion driven processes. We next use our system to explore how molecular crowding might influence clustering in Fig. 13. We employ 320 spherical crowders of radius = 5 nm (Fig. 13B). We see a large increase in average cluster size and occupancy upon adding the crowder. This enhancement of clustering can be attributed to an excluded volume effect, which increases the effective concentration (or thermodynamic activity) of the reactive molecules by reducing the available reaction volume. Although a simple calculation shows that our spheres fill ~1/6 of the 10^6^nm^3^ reaction volume, the actual excluded volume might be better approximated by subtracting the packed volume of the spheres from the total volume; if we assume a cubic packing, this volume is 320 X (2r)^3^, or ~1/3 of the 10^6^nm^3^ simulation volume. To test this effect of volume reduction, we modeled an additional case where 36 interacting molecules are put into a 2/3 of reference reaction volume with no crowders (Fig. 13C). The increase in average cluster size or occupancy is less than the crowded system. So the extent of volume exclusion is higher than the sum of closely packed volume of the crowder. This is likely a consequence of an additional excluded volume contribution from the finite size of the sites within the 36 interacting molecules. That is, the mass-center of each interacting molecule can’t access a spherical volume corresponding to the sum of its own radius and that of the crowder (Fig. S8). Also noteworthy is the slower kinetics of cluster formation in the crowded environment Fig. 10B, inset). This cannot be attributed simply to slower diffusion, as the diffusion of a single NWASP is not strongly retarded (Fig. S9). These results emphasize the importance of properly accounting for the sizes of macromolecules when modeling reaction networks containing multivalent interactions.

**Figure 13:**
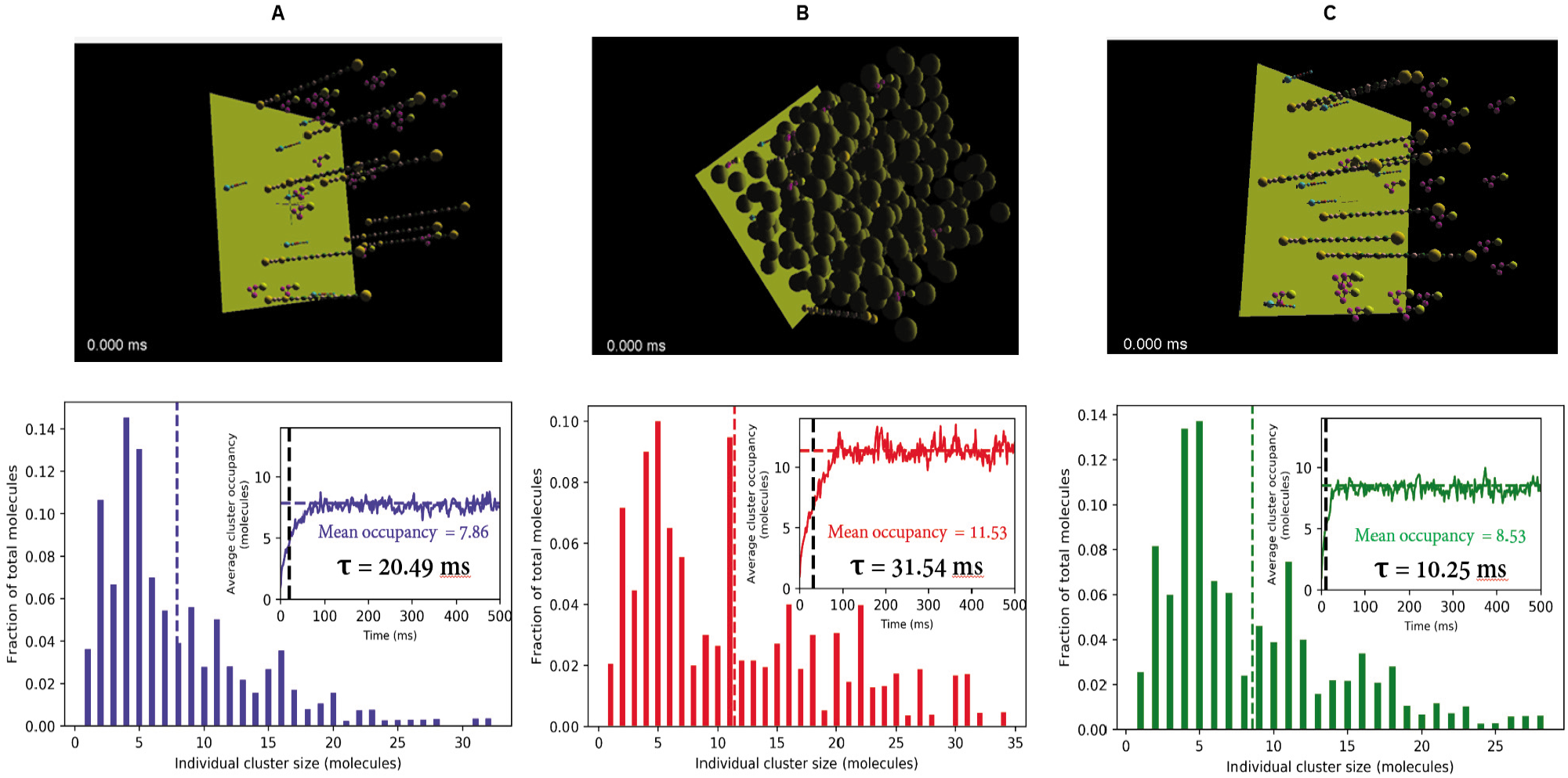
Molecular crowding enhances clustering beyond a simple excluded volume concentration effect. (A) Reference system with 36 interacting molecules in a 100^3^ nm^3^ cubic reaction volume. (B) System with 320 inert crowder (radius = 5 nm) along with 36 interacting molecules in the reference reaction volume. The crowders would take up ~1/3 of the volume based on cubic packing (each sphere with radius r would occupy an effective cubic volume having a side length of 2r). (C) 36 interacting molecules in 2/3 of the reference volume, i.e., (100*100*67) nm^3^. The Corresponding steady state distributions of molecular occupancies are shown in the right hand panels (inset shows the time course; τ is the characteristic time representing how fast the system converges to steady state). Since the crowded system takes longer to reach steady state, four time points (*50 runs = 200 realizations) are sampled for the distribution.

## Discussion

The aims of our study were to establish the biophysical factors that shape how weakly binding multivalent interactions control the formation of dynamic clusters, which have been also called molecular ensembles. These are broadly important because molecular ensembles can serve as functional platforms for key cellular signaling systems. We have developed a kinetic model that utilizes a coarse representation (Figs. 1 and S1) of the key molecular features in 3 interacting multivalent molecules. It is inspired by the nephrin/Nck/NWASP system that underlies the structural integrity of the kidney filtration unit (33, 34). The multivalent binding domains from each of these molecules also served as the basis for a seminal study of microdroplet formation by liquid-liquid phase separation (23). It is important to emphasize that our model is not intended to quantitatively predict the ability of this system to phase separate. This would require us to account for weak non-specific interactions (e.g. electrostatic) (17, 35). However, cluster formation through multivalent interactions is a definite prerequisite for liquid droplet phases, and if we push our own system to high concentrations we see behavior that has the hallmarks of phase separation (Fig. 6). Our study primarily focusses on elucidating the molecular and cellular structural features that control cluster formation and size.

Our reference system, which is below the saturation boundary for phase separation, consisted of 6 membrane-bound nephrin molecules, 20 cytosolic Nck molecules and 10 cytosolic NWASP molecules in a cubic reaction volume of 100nm on a side; one face of the cube represented a planar patch of membrane within which the nephrin molecules could diffuse (Fig. 2 and Table 1). This stoichiometry was close to the optimal for maximizing interaction between all the binding sites on these multivalent molecules. Based on stochastic simulations, we analyzed the kinetic approach to steady state, the distribution of cluster sizes at steady state and the distribution of molecular occupancies within each possible cluster size (Fig. 3). We found the reference system of 36 molecules was sufficiently large to simulate all the key features of the system (Fig. 4) because, perhaps somewhat surprisingly, the cluster size and cluster occupancy histograms were very similar for 18, 36 or 144 molecules. This gave us confidence that the reference system properties could serve as a good baseline for computational experiments that systematically probed for the effects of molecular and cellular structure on the clustering behavior. It also demonstrated that a balance of enthalpic and entropic factors limited the size distribution of clusters and prevented their annealing into a single large complex.

As would be expected, decreasing valency decreases the steady state cluster size distribution (Fig. 7). This computational experiment mirrors the trend in an in vitro experiment by the Rosen lab (23), in which nephrin constructs with 3, 2 and 1 phosphotyrosines required progressively higher concentrations of Nck and NWASP to produce phase separation. But, more subtly, the simulations show that there are preferred cluster sizes with specific monomer compositions that are dependent on valency, with lower valency sharpening the preference for clusters of optimal stoichiometric composition. This behavior is accentuated when the binding affinity between sites is increased (Fig. 5) and can be attributed to stoichiometry matching, whereby clusters may become limited to stoichiometries where all binding sites are fully occupied. A similar effect was shown to be able to control the size of clusters in a binary system, consisting of an octavalent and tetravalent binding pair, to model the membraneless pyrenoid organelle in chloroplasts (30). When the tetravalent molecule was altered to a trivalent molecule, the cluster size increased despite the reduction in valency.

We deconvolved the influence of steric interactions, stoichiometry matching and molecular flexibility on steady state cluster sizes by systematically altering NWASP structural features (Figs. 8, 9, 10). In SpringSaLaD, each spherical site excludes volume to represent steric effects and also serves as a pivot to impart molecular flexibility; the stiff spring links serve to maintain a fixed distance between the sites and also transmit forces. Decreasing flexibility by removing sites that were not involved in binding produced a major increase in cluster sizes. We showed that this effect is not due to a decrease in local steric interference to binding, which might also be an effect of removing these structural sites (compare Fig. 10A and C). We believe this effect can be attributed to a loss in entropy during cluster formation, where more flexible monomers would lose more entropy than less flexible monomers. Another way of thinking about this is to consider that the less flexible molecules would tend to adopt more open conformations with more exposed binding sites. Stoichiometry matching is also controlled by the geometric spacing between the respective binding sites; if the sites are separated by rigid linkers, only those multivalent interactions with matching spacing between binding pairs can produce ladder-like linkages to limit cluster size (Figs. 10B vs. 10E and 10D vs. 10F). If the distances between binding sites are more flexible (Figs. 10A and 10C), stoichiometry matching between multivalent pairs can again restrict the growth of larger clusters.

We examined steric interactions by changing the sizes or eliminating the structural sites in NWASP, keeping the sizes of binding sites constant (Figs. 9 and S5). The most surprising conclusion was the dominance of the peripheral structural sites in NWASP; removing or shrinking these sites significantly increased cluster sizes (Fig. 9). Shrinking the structural sites that were located between binding sites without changing the peripheral sites (Fig. 10C) actually slightly shifted the cluster occupancy histogram to smaller sizes. Apparently, the crosslinking of binding sites favors exclusion of the structural sites from the interior of clusters, with larger structural sites having a greater propensity for exclusion (Fig. S6). This self-organizing effect would lower the entropy and explain the greater steric effect for peripheral domains. As an alternate view, these large structural sites at the periphery of clusters would serve as a steric barrier to the recruitment of additional monomers to binding sites in the interior, thereby limiting cluster expansion. A biological implication is that these domains would be present at the exterior where they could participate in downstream signaling. In particular, the VCA domains of NWASP (within the right-most yellow structural site in Fig. 1C. 2) would be assembled at the periphery of clusters; this is particularly advantageous, because a pair of proximate VCA domains is required to recruit and activate ARP2/3, which in turn, nucleates branched actin polymerization (36, 37). In general, such non-linearity is a hallmark of signaling systems.

Turning to cellular effects, our model shows increased steady state clustering in the presence of inert crowders (Fig. 13). We attribute this to an effective increase in binding site concentration due to the excluded volume of the spherical crowders. On the other hand, crowders significantly reduce the rate of cluster formation. A similar, but subtler consideration of effective concentration can explain how membrane anchoring of nephrin significantly promotes clustering (Fig. 11). This is because confining reactions to the membrane effectively reduces the available reaction volume once the initial complement of Nck and NWASP molecules are recruited from the bulk volume. Thus, annealing of small membrane-associated clusters into larger ones is favored because the binding sites are at an effectively higher concentration. This effect is further enhanced when the surface area of the membrane is decreased while keeping the bulk concentrations constant (Fig. 12): increasing the membrane density 4-fold doubles the average cluster size occupancy. Lipid rafts provide a biological platform to concentrate membrane proteins and are thought to play an especially important role in receptor-mediated signaling (38). Our results suggest that clustering can serve as a positive feedback mechanism to amplify the ability of lipid rafts to localize membrane receptors and thereby amplify spatially encoded signals.

Our computational experiments have allowed us to gain insights into the biophysical features that control the formation of molecular ensembles. But most importantly, they suggest experiments that will help to validate these ideas. In vitro experiments employing manipulated protein constructs and/or supported membranes, similar to the work from the Rosen lab (39, 40), could be used to systematically test our predictions on how steric hindrance, linker flexibility and membrane surface density influence the formation of molecular ensembles. Single molecule or super resolution microscopy experiments would be especially pertinent for characterization of the size and composition of molecular ensembles in cells. It would be of great interest to explore our prediction, for example, that the VCA domain of NWASP is most likely to be situated at the periphery of clusters. Also, disrupting rafts through cholesterol depletion prior to stimulation by ligand, can test our general prediction that confinement of receptors to lipid rafts will increase the size of clusters. Ultimately, we anticipate that coarse-grained molecular kinetic modeling can guide experimental manipulation of molecular ensembles to control downstream cellular responses.

## Supporting information

Supplemental movie of cluster formation

## Acknowledgments

We gratefully acknowledge useful discussions with Michael Blinov, James Schaff, Boris Slepchenko and Paul Michalski. We are grateful to the referees for their thorough and insightful comments on the first version of this paper. The research was supported by National Institute of General Medical Science grant P41 GM103313.

## Author Contributions

AC performed research, designed statistical analyses, analyzed data and co-wrote the manuscript. MY performed initial pilot studies. LL conceived the project, performed research and co-wrote the manuscript.

