## Supplemental Figures and Tables for "The interplay of structural and cellular biophysics controls clustering of multivalent molecules"

**Aniruddha Chattaraj, Madeleine Youngstrom and Leslie M. Loew***

**Supplementary Materials**

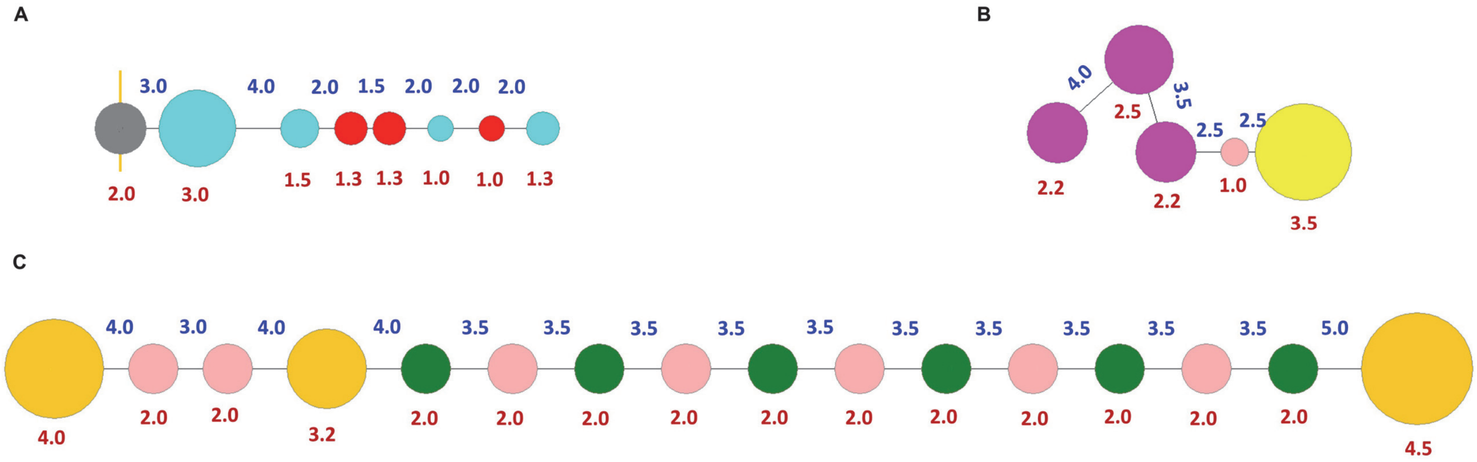


**Figure S1: Structural details of model components**. (A) nephrin, (B) Nck, (C) NWASP. Diameter of the sites are shown in brown below the corresponding site. Center-to-center distance between sites are shown in blue. All distance units are nm.

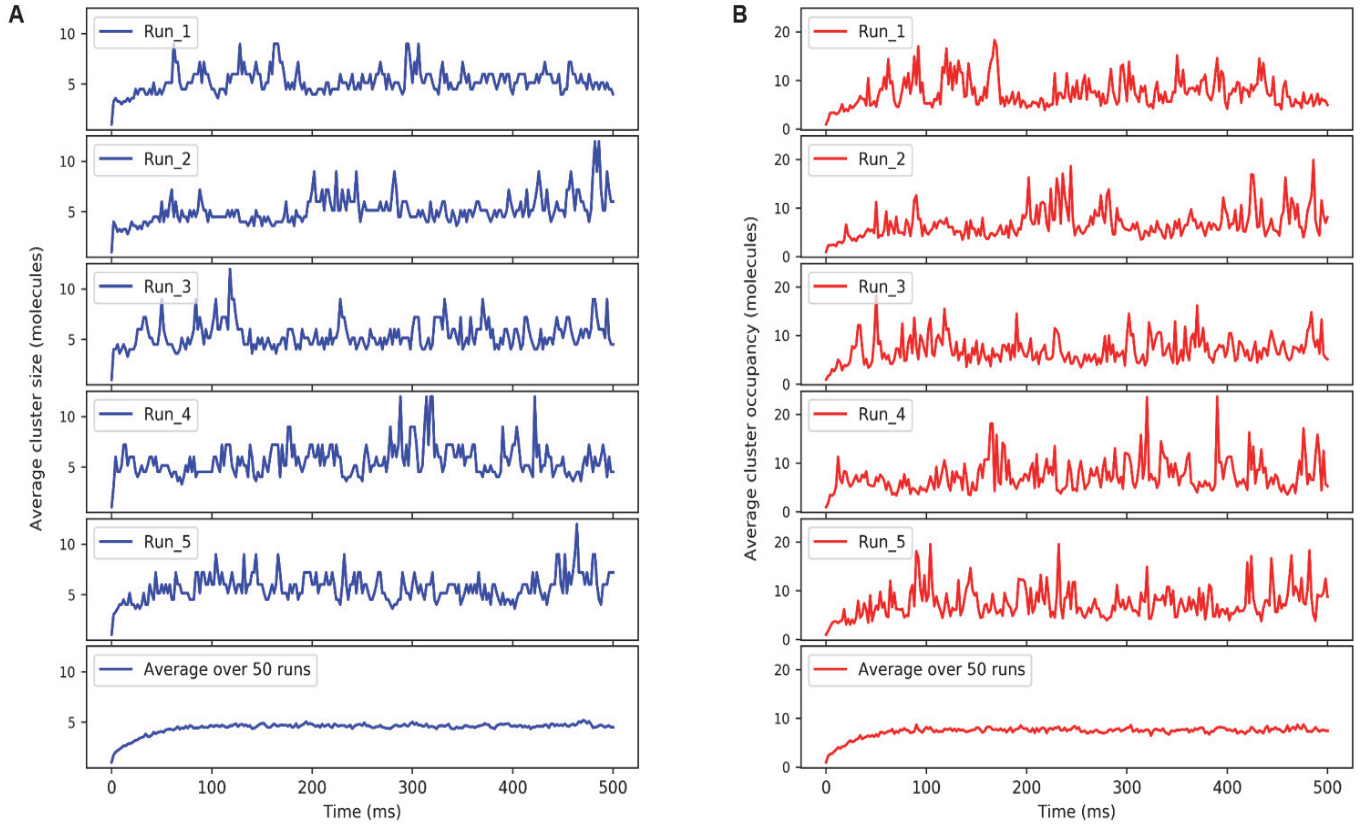


**Figure S2: A sampling of 5 individual trajectories.** (A) Average cluster size, (B) Average cluster occupancy. The corresponding average over 50 trajectories are at the bottom.

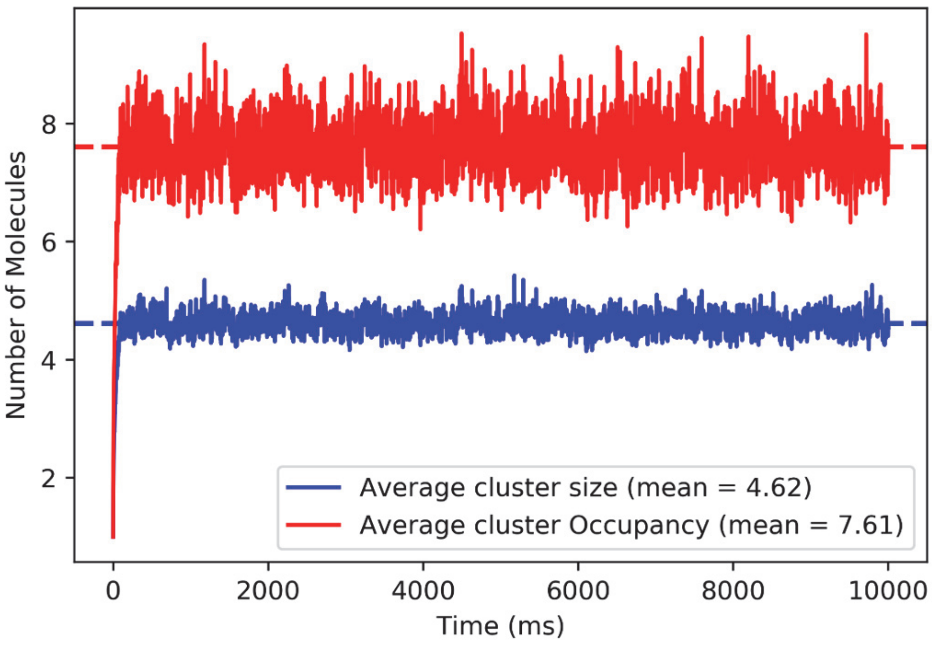


**Figure S3: Average cluster size for the reference system is stable over 10s**. Red and blue trajectory represents the molecular occupancy and average cluster size respectively.

### (A) 2X concentration

| System | X (nm) | Y (nm) | Z (nm) | # molecules |
|---|---|---|---|---|
| Vol_1x | 50 | 100 | 100 | 36 |
| Vol_2x | 100 | 100 | 100 | 72 |
| Vol_4x | 200 | 100 | 100 | 144 |

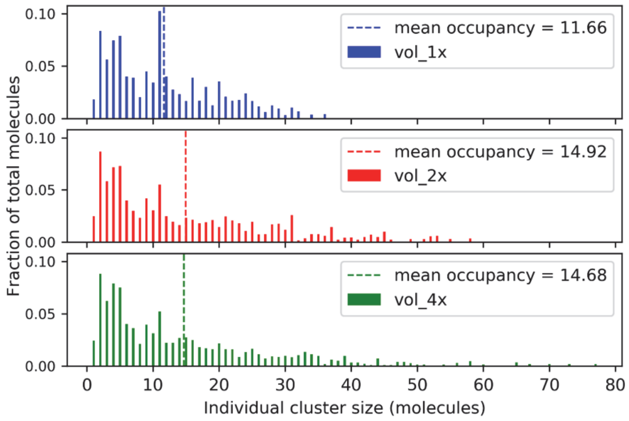


### (B) 4X concentration

| System | X (nm) | Y (nm) | Z (nm) | # molecules |
|---|---|---|---|---|
| Vol_1x | 50 | 50 | 100 | 36 |
| Vol_2x | 50 | 100 | 100 | 72 |
| Vol_4x | 100 | 100 | 100 | 144 |
| Vol_8x | 200 | 100 | 100 | 288 |
| Vol_16x | 200 | 200 | 100 | 576 |

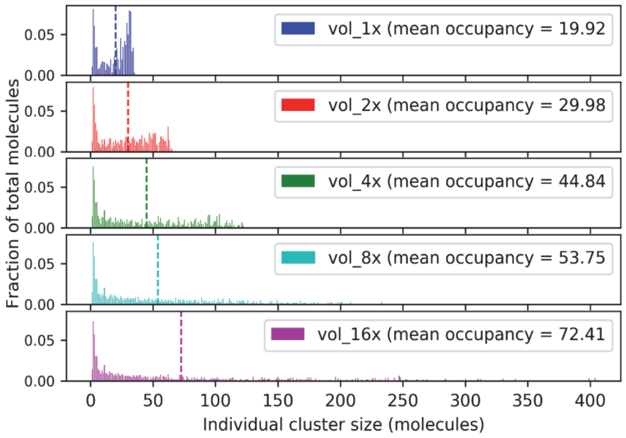


**Figure S4: System size dependence at increased concentration**. Molecular concentrations are increased by a factor of (A) 2 and (B) 4, relative to the reference system (Table 1)

**Table S1: Stoichiometric composition of individual clusters**

| System | Cluster size | Major composition (nephrin, Nck, NWASP) | Occurrence |
|---|---|---|---|
| Reference | 2 | 0, 1, 1 | 96 % |
| | 4 | 1, 2, 1 | 98 % |
| | 5 | 1, 3, 1 | 96 % |
| | 6 | 1, 3, 2 | 91 % |
| | 9 | 2, 5, 2 | 96 % |
| | 11 | 2, 6, 3 | 92 % |
| Bivalent neprin | 4 | 1, 2, 1 | 99 % |
| Monovalent nephrin | 5 | 2, 2, 1 | 98% |

### Table S2: Effect of initial stoichiometry on cluster size

| Molecular Stoichiometry (nephrin, Nck, NWASP) | Mean cluster size (molecules) | Mean Cluster occupancy (molecules) |
|---|---|---|
| 6, 20, 10 (Reference) | 4.71 | 7.86 |
| 6, 10, 20 | 1.90 | 3.43 |
| 10, 20, 6 | 4.70 | 7.33 |
| 10, 6, 20 | 1.45 | 2.29 |
| 20, 6, 10 | 1.46 | 2.39 |
| 20, 10, 6 | 1.98 | 3.77 |
| 12, 12, 12 | 2.49 | 4.56 |
| 7, 22, 7 | 4.48 | 7.78 |
| 7, 18, 11 | 4.42 | 7.08 |
| 8, 22, 6 | 4.57 | 7.79 |
| 5, 22, 9 | 3.79 | 6.61 |
| 4, 24, 8 | 2.96 | 5.41 |

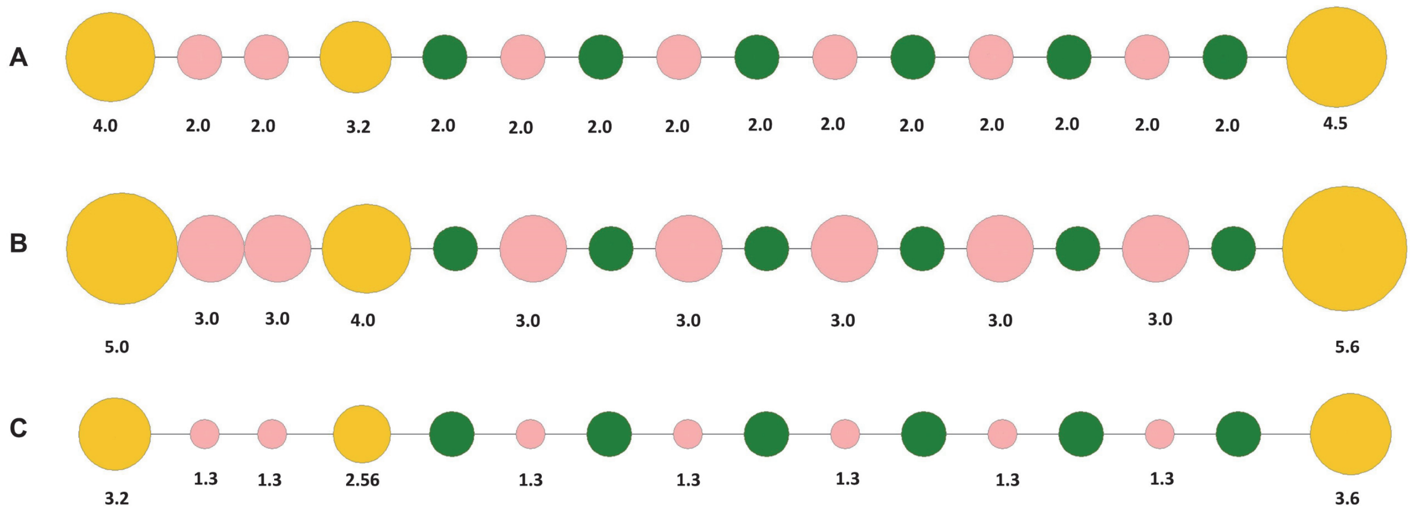


**Figure S5: Structural details of NWASP configurations used for steric test.** (A) Reference (B) Larger structural sites (C) Smaller structural sites. Diameter of all the sites are shown for reference structure. For B & C, only those sites are shown whose diameters are varied; rest of sites retain same sizes as reference configuration.

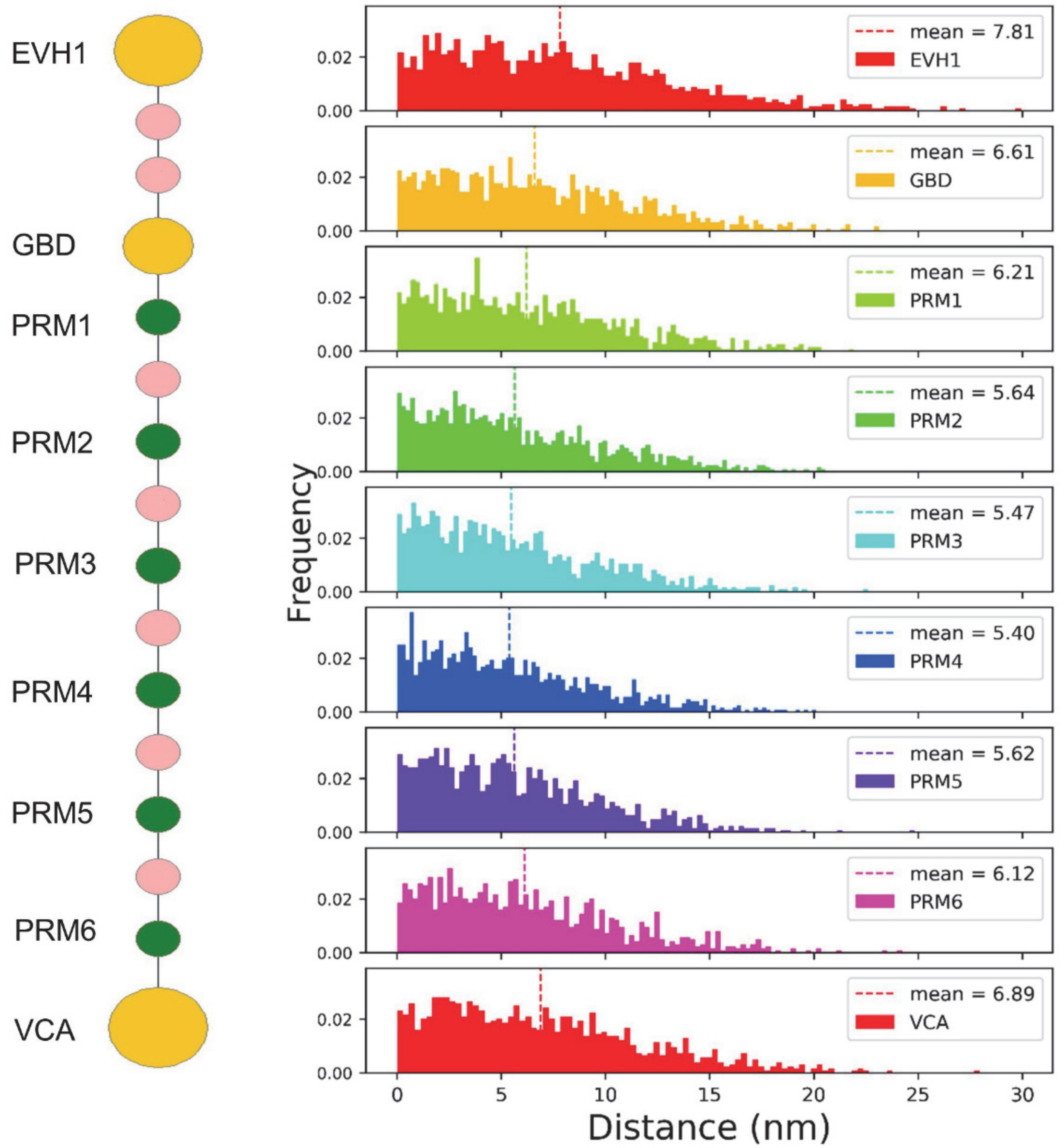


**Figure S6. Peripheral sites of NWASP are located at the periphery of large clusters.** Distance of different NWASP sites from the centroid of the respective cluster (analyzed for clusters size >= 10 molecules)

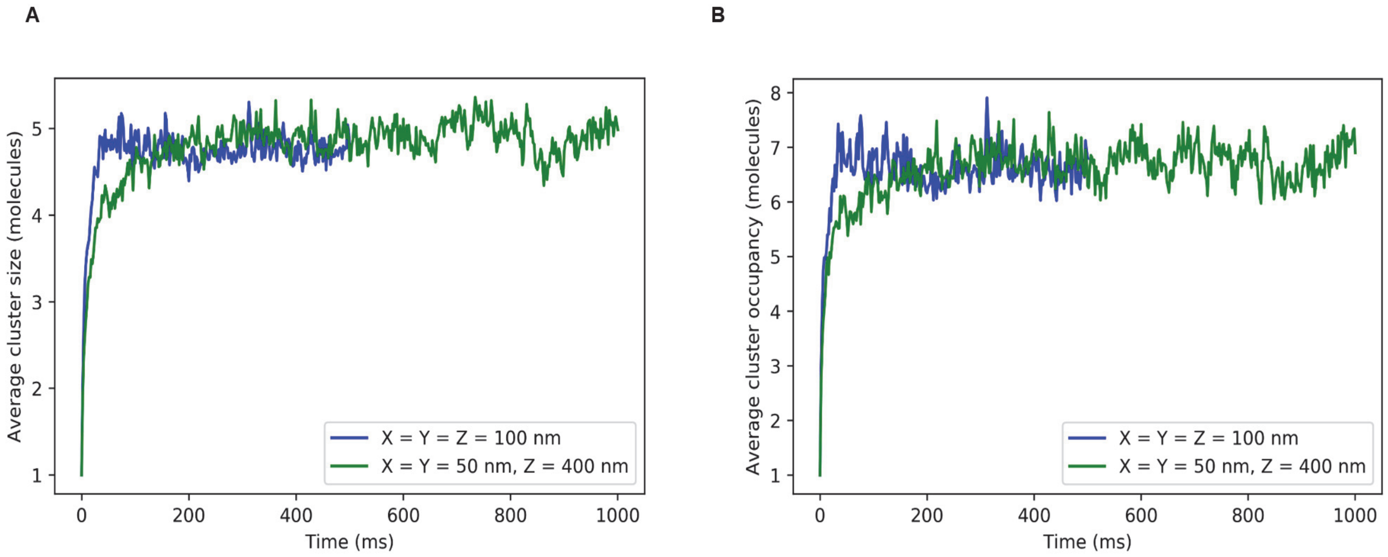


**Figure S7: In the absence of membrane, change in volume aspect-ratio does not alter the steady state cluster size or occupancy, but does alter the kinetics.** Dynamics of average (A) cluster size and (B) occupancy

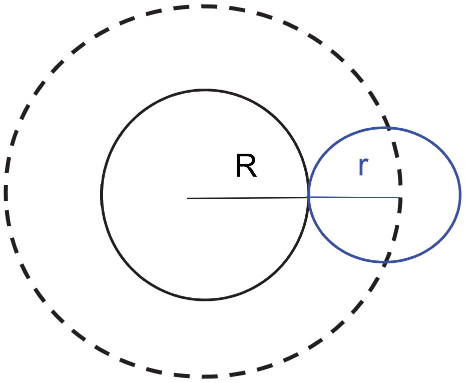


**Figure S8: Excluded volume effect.** Blue and black (solid) circle represents the biochemically active (test) and inert (crowder) molecules respectively with radius r and R. Due to its finite size, the test molecule can't access an additional volume (shown in dotted black circle) around the crowder. So the excluded volume with the addition of crowder molecules is greater than the sum of their close-packed volume.

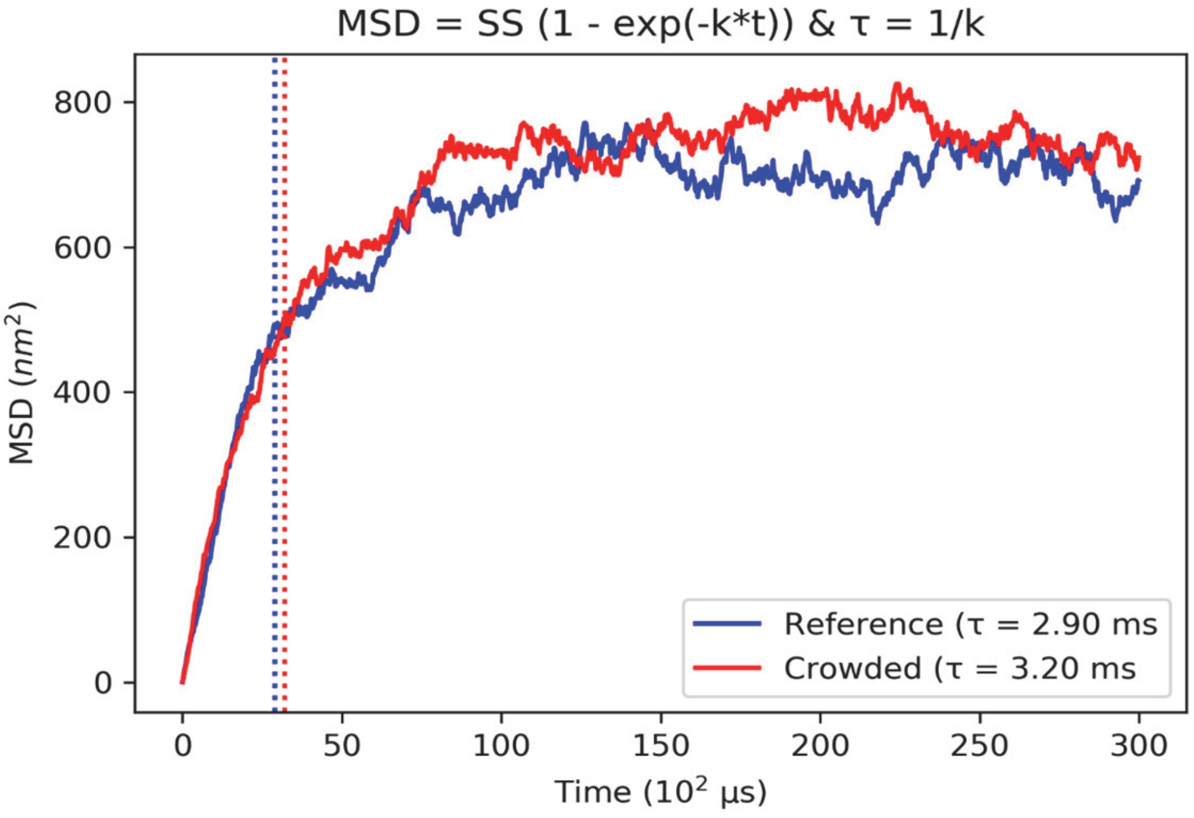


**Figure S9. The presence of crowders has a minor effect on the diffusion of a single NWASP molecule.** The diffusion of the molecular centroid is monitored. Blue, reference environment (in presence of 6 Nephrin, 20 Nck, 9 NWASP). Red, Crowded environment (reference + 320 inert molecule, diameter 5 nm). The binding interactions were turned off for these simulations.
